# Targeting targeted memory reactivation: characteristics of cued reactivation in sleep

**DOI:** 10.1101/2021.12.09.471945

**Authors:** Mahmoud E. A. Abdellahi, Anne C. M. Koopman, Matthias S. Treder, Penelope A. Lewis

## Abstract

Targeted memory reactivation (TMR) is a technique in which sensory cues associated with memories during wake are used to trigger memory reactivation during subsequent sleep. The characteristics of such cued reactivation, and the optimal placement of TMR cues, remain to be determined. We built an EEG classification pipeline that discriminated reactivation of right- and left-handed movements and found that cues which fall on the up-going transition of the slow oscillation (SO) are more likely to elicit a classifiable reactivation. We also used a novel machine learning pipeline to predict the likelihood of eliciting a classifiable reactivation after each TMR cue using the presence of spindles and features of SOs. Finally, we found that reactivations occurred either immediately after the cue or one second later. These findings greatly extend our understanding of memory reactivation and pave the way for development of wearable technologies to efficiently enhance memory through cueing in sleep.

## Introduction

Memories are neurally replayed during sleep, and this process is associated with consolidation^1–3^. Targeted memory reactivation (TMR) is a technique in which sensory cues are paired with learned material during wake, then re-presented during subsequent sleep in order to trigger reactivation of the associated material^4,5^. This procedure leads to memory benefits for reactivated material in Non-REM (NREM) sleep (see^5^ for a recent meta-analysis). Importantly, several studies have confirmed the reinstatement of learning related brain activity after TMR cues in NREM sleep (see^6^ for a review). Studies have looked at the neural structures involved in reactivation^7,8^, and found both positive^8–11^, and negative^12^ relationships between the extent of reactivation and subsequent memory benefits.

Cortical activity during slow wave sleep (SWS) is characterised by high amplitude slow oscillations (SOs) in which neurones oscillate between hyperpolarization with neuronal silence (“down-state”) and depolarisation with sustained firing (“up-state”). Depolarised SO up-states drive memory reactivation in the hippocampus via interactions with thalamic sleep spindles (SS) and hippocampal sharp wave ripples (SWRs)^13^. Studies have shown that stimulation during the up-going SO phase is associated with greater memory benefit compared to the down-going phase^14,15^, and that stimulating the up-going phase produces a higher ERP response compared to f the down-going phase^16^. This could be due to the fact that neurones are in the process of depolarising and are thus moving closer to the threshold for firing during the up-going phase. Furthermore, fast spindles, which have been linked to both memory consolidation^17^ and reactivation^9^, typically occur on the up-going phase^18,19^.

TMR is thought to prime a memory trace for reactivation^6^, and has been shown to trigger SO-spindle complexes^9,20,21^. We predict that application of such priming during the up-going phase of the SO just prior to a spindle event may lead to reactivation. On the other hand, application of stimulation during the down-going phase of the oscillation when fast spindles rarely occur and excitability is reduced, is less likely to produce reactivation.

SOs vary in terms of generation locus as well as shape, for instance having different periods, trough depths, and peak to trough slopes. These varied morphologies are thought to relate to the degree of synchronisation across neural populations in the cortex^19,22^. Given these differences, some SOs are likely to facilitate reactivation more efficiently than others. We hypothesise that it may be possible to predict this efficiency based on features of the ongoing oscillatory structure of sleep, with specific reference to SOs and spindles, in the time period directly before stimulation. This would not only optimise stimulation, but also allow more targeted stimulation, minimising the number of sound cues needed to influence consolidation.

In the current work, we set out to characterise memory reactivation after TMR in NREM sleep and to determine whether TMR on the up-going phase is more likely to elicit reactivation compared to the down-going phase. Additionally, we ask whether we can predict the optimal time for TMR stimulation using ongoing morphology of SOs and spindles. We use a serial reaction time task (SRTT). In the SRTT, participants respond to audio-visual cues by pressing four buttons, one per finger, with two fingers on each hand (Fig. 1). Each press was cued by a picture-sound pair, and tones associated with the task were replayed during SWS on the post-training night to elicit memory reactivation. Importantly, prior to performing the task, participants were exposed to the tones during an adaptation night. The addition of this adaptation night provided control data during which tones could not have evoked memory reactivation, as they were not yet linked to the task. We then trained a classifier to identify neural responses associated with left and right-handed presses in wake, and applied it after each TMR tone in SWS on both adaptation and experimental nights. We also used the features of the ongoing oscillation to train another classifier to determine whether TMR applied at a given time in the oscillatory sequence would elicit detectable reactivation.

**Fig. 1.**
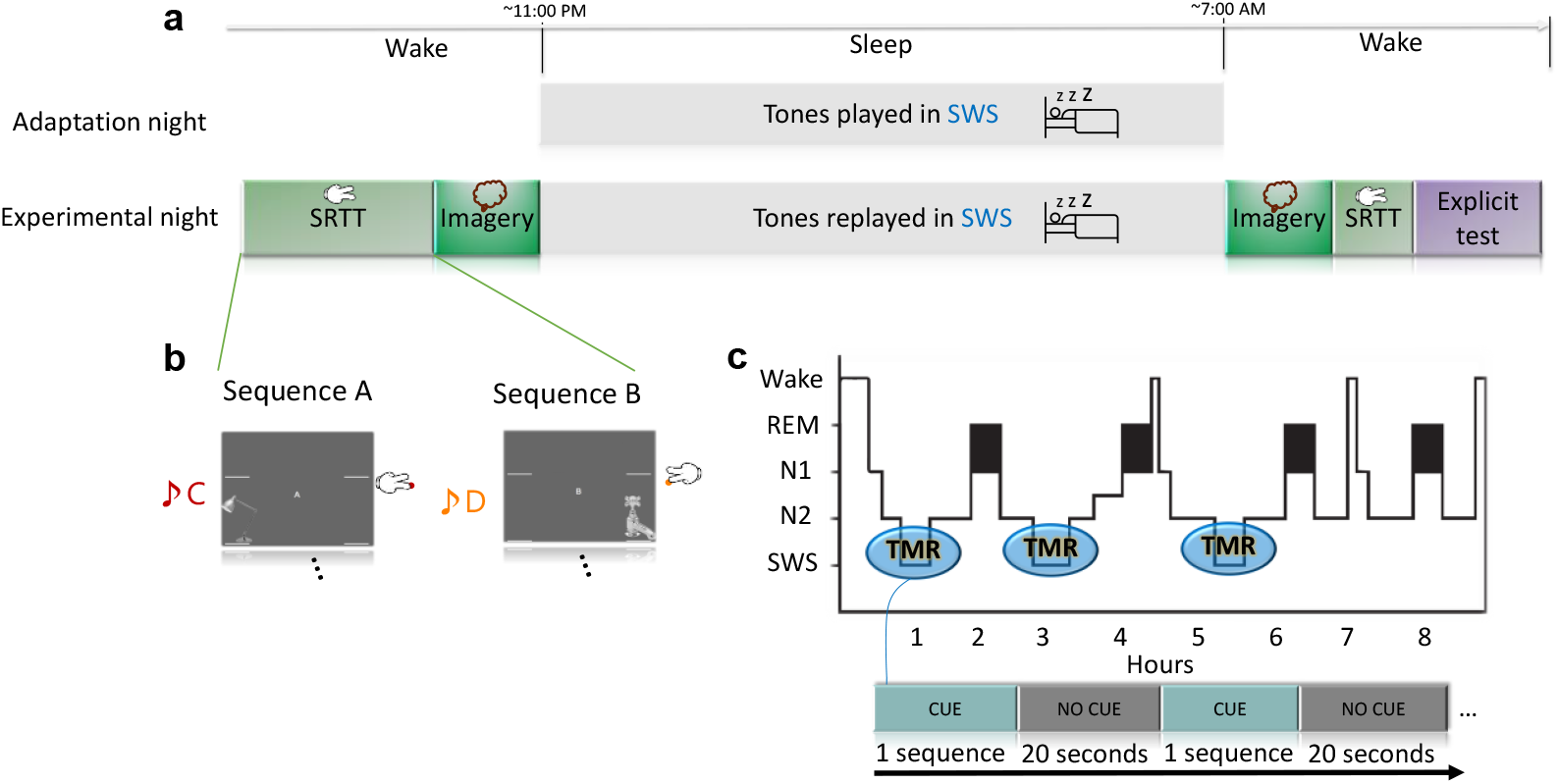
Experimental design. **a** Participants spent two nights in the lab, Firstly, an adaptation night during which EEG recordings were acquired and tones were presented to the participants during SWS. Secondly, in the experimental night, participants completed the serial reaction time task (SRTT), followed by motor imagery task. In the imagery task, the participants were cued with pictures and sounds, but were told to only imagine performing the finger tapping (without movement). Afterwards, participants slept in the lab and TMR cues were presented during SWS. After waking up, participants completed the motor imagery and then SRTT, and finally an explicit recall task in which they marked the locations of the images as they had appeared in the sequences as accurate as they could. **b** In the SRTT, images are presented in two different sequences each with a different set of tones. Each image is associated with a unique tone and requires a specific button press. In the imagery task, participants hear the tones and see the images as in SRTT but only imagine pressing the buttons. **c** The sounds of only one learned sequence (cued sequence) were re-played during SWS sleep, with a 20 second pause between repetitions.

## Results and Discussion

### TMR improved sequence memory

Prior work has shown that SRTT is facilitated by TMR in SWS^23–25^. Our data are in line with this, since cueing led to significant overnight behavioural improvement for the cued sequence compared to non-cued (t-test, n=15, p = 0.042), see^26^ for full behavioural analysis.

### Multiple reactivations detected after TMR

Prior work^9,10^ has suggested a recurrent pattern of reactivation after a TMR cue, with a reinstatement of the target memory immediately after the cued memory followed by a later reinstatement, see^6^ for a discussion. Building on this work, we examined the time course of classification after TMR for evidence of a similar pattern. Our results revealed significantly higher classification performance in the experimental night than the adaptation night, with two different effects described by two clusters after TMR onset (Fig. 2a). An early cluster (p = 0.02) that occurred immediately after TMR onset and a late cluster (p = 0.01) that occurred ~1sec later. Results are corrected for multiple comparisons with cluster-based permutation (see methods for details), trial duration in sleep was 1,500ms.

**Fig. 2.**
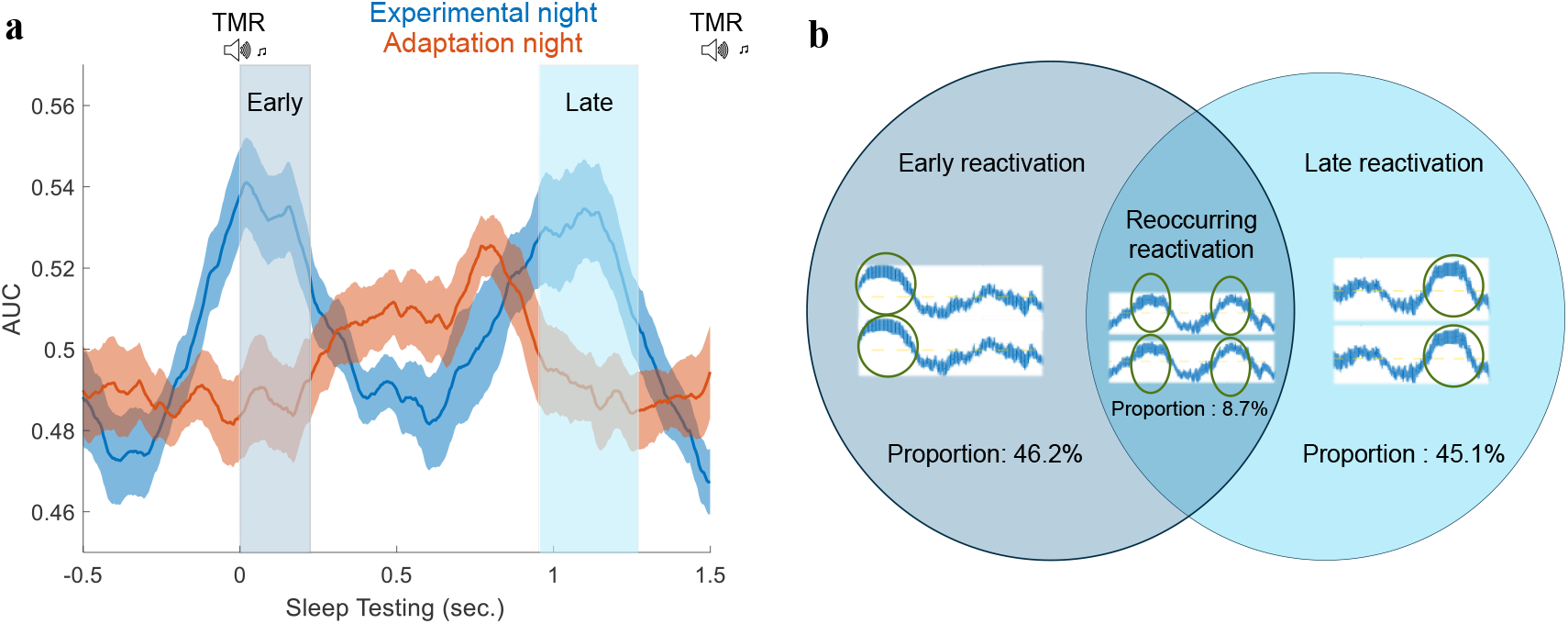
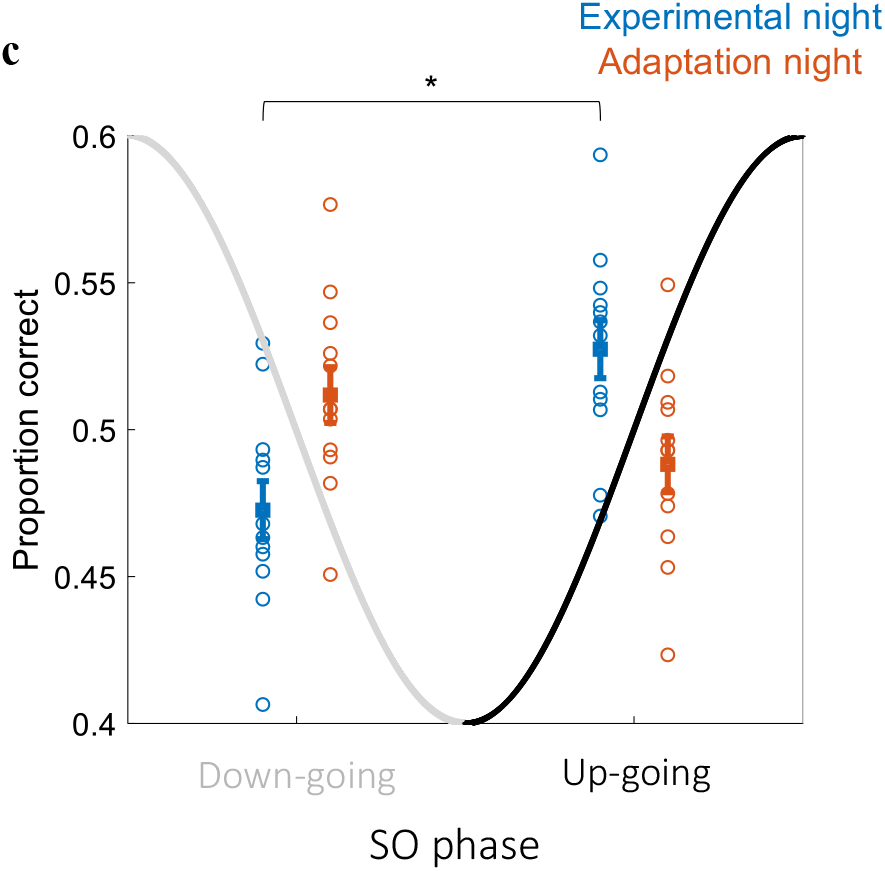
Detectable reactivations and their characteristics. **a** Classification results for both nights. The blue curve represents the area under the ROC curve (AUC) across time for the experimental night (with the mean represented by a solid curve and standard error shaded around it), red curve represents results of the adaptation night. TMR sounds were presented at the beginning of sleep trials, ‘early’ and ‘late’ are used to mark early and late reactivations. Classification results have two significant effects expressed by two clusters, (early cluster, p = 0.02, and late cluster, p = 0.01). **b** Proportions of correct trials with only early reactivation (46.2%), only late reactivation (45.1%), and reoccurring reactivations (8.7%). **c** Proportion of correct trials with the TMR cue falling on different SO phase transitions for the two nights. Each circle represent data from one participant and the curve is a simplified cartoon representation of the phase of a SO, the two SO phases are marked on the x-axis and the y-axis represents the proportion of correct trials. The preferred phase for early reactivation is when the sound falls on the up-going transition of the SO (Wilcoxon signed rank test, n=12, P = 0.019, Z = 2.4) compared to down-going.

To test whether this was due to recurrent reactivation of the same response, we examined each trial to see whether it included an early reactivation, a late reactivation, or both. We then looked at whether the same trials were classified correctly at both early and late peaks (Fig. 2b). This revealed that the majority of trials contain one reactivation, either early or late, and only 8.7% of trials showed reoccurring reactivation by classifying correctly during both early and late peaks. Comparison of reoccurring reactivation to chance level showed that it was below chance (Wilcoxon signed rank test, n = 12, p = 0.002, z = −3.0594) (see methods for details); (Fig. 2b).

Overall, these results suggest that the reactivations we are detecting are not recurrent, but instead commonly occur just once after each cue: either early or late within our trial duration. It is possible that this may also have been the case in the prior human studies^9,10^, as they reported the performance across many trials and did not examine individual trials.

### Preferred TMR phase for reactivation

There is evidence that TMR may be more effective when applied to the up-going phase of the SO^14,27,28^. Moreover, fast rhythms, such as spindle, and gamma activity are more prominent in the SO up-going state than in the SO down-going state^29–31^, also there are changes to the ERP when the auditory stimulation is applied during the up-going phase of the SO^16^. Building on the extensive literature relating to reactivation during rodent sharp-wave ripples^32–34^, data from human epilepsy patients has shown that the SO up-going state shows more sharp-wave ripples and gamma oscillations^35^.Sharp-wave ripples have been shown to carry reactivation^36^, on the other hand, they are supressed during the SO down-going state^37^. Thus, the up-going SO phase appears to be the preferred time for stimulation to improve memory^14^.

Given this background, we predicted that TMR would more effectively trigger reactivations if applied to the up-going phase of the SO. We tested this by dividing correctly classified sleep trials based upon the phase at which TMR was initiated, see methods for details. In the experimental night, this showed a significantly higher proportion of correct trials for early reactivation when TMR was applied on the up-going compared to the down-going SO transition and chance level of 0.5 (Wilcoxon signed rank test, n=12, p = 0.019, z = 2.4), (Fig. 2c). As a control, we compared the proportion of correct trials from the adaptation night for up-going and down-going SO phases and chance level (Wilcoxon signed rank test, n = 12, p = 0.24, z = 1.18). We repeated this analysis for the incorrectly classified trials for early reactivation and found no significant difference between transitions and chance level for either experimental night (Wilcoxon signed rank test, n = 12, p = 0.16, z = 1.4) or adaptation night (Wilcoxon signed rank test, n = 12, p = 0.31, z = 1.02). We also did the same for late reactivation but found no difference between up-going and down-going phase transitions (p > 0.2).

This analysis shows that TMR cues which fall on the up-going transition of the SO, are more likely to lead to a classifiable early reactivation, than TMR cues that fall on the down-going phase supporting the idea that SOs interact with reactivation in some functional way. This could also be important for optimisation of TMR cueing in order to successfully trigger reactivation.

### Predicting reactivation using pre-cue SO features

While the literature suggest that reactivation is modulated by SOs^1,38^, the mechanism for this modulation remains to be understood. We were interested to determine whether the features of the ongoing SO prior to stimulation, could predict whether a given TMR cue would produce a classifiable reactivation. In other words, we wanted to know whether some points in the oscillatory pattern are more optimal than others for delivering TMR, and if so, which features of the ongoing oscillatory structure determine this. To examine this, we performed a second classification analysis, this time training our classifier on pre-cue SO features. We wanted to see if we could discriminate between correct and incorrect trials (based on the results of main reactivation classifier, Fig. 2a). To this end, we extracted SO features from Fz electrode during the two seconds of data before the onset of TMR.

The extracted features are described in Supplementary Table 1. These features were fed to decision tree classifiers^39^ which were trained on two classes: correctly classified, and incorrectly classified from the main classifier, see methods for details. As a control, we compared the results obtained from the SO-based classifier of the experimental night to the SO-based classifier trained and tested using the adaptation night (Fig. 3a). The performance of the experimental night classifier was significantly higher than that of the adaptation night for predicting the early reactivation (Wilcoxon signed rank test, n=12, p = 0.015, z = 2.43), but not for predicting the late reactivation (p > 0.2). This indicates that it is possible to predict classifiable early reactivation in the experimental night when learned information could actually be reactivated, compared to the control condition when the task had not been learned yet. This result shows that we can use SO features to predict the optimal time to deliver TMR, in order to maximise the probability of producing a classifiable early reactivation.

**Fig. 3.**
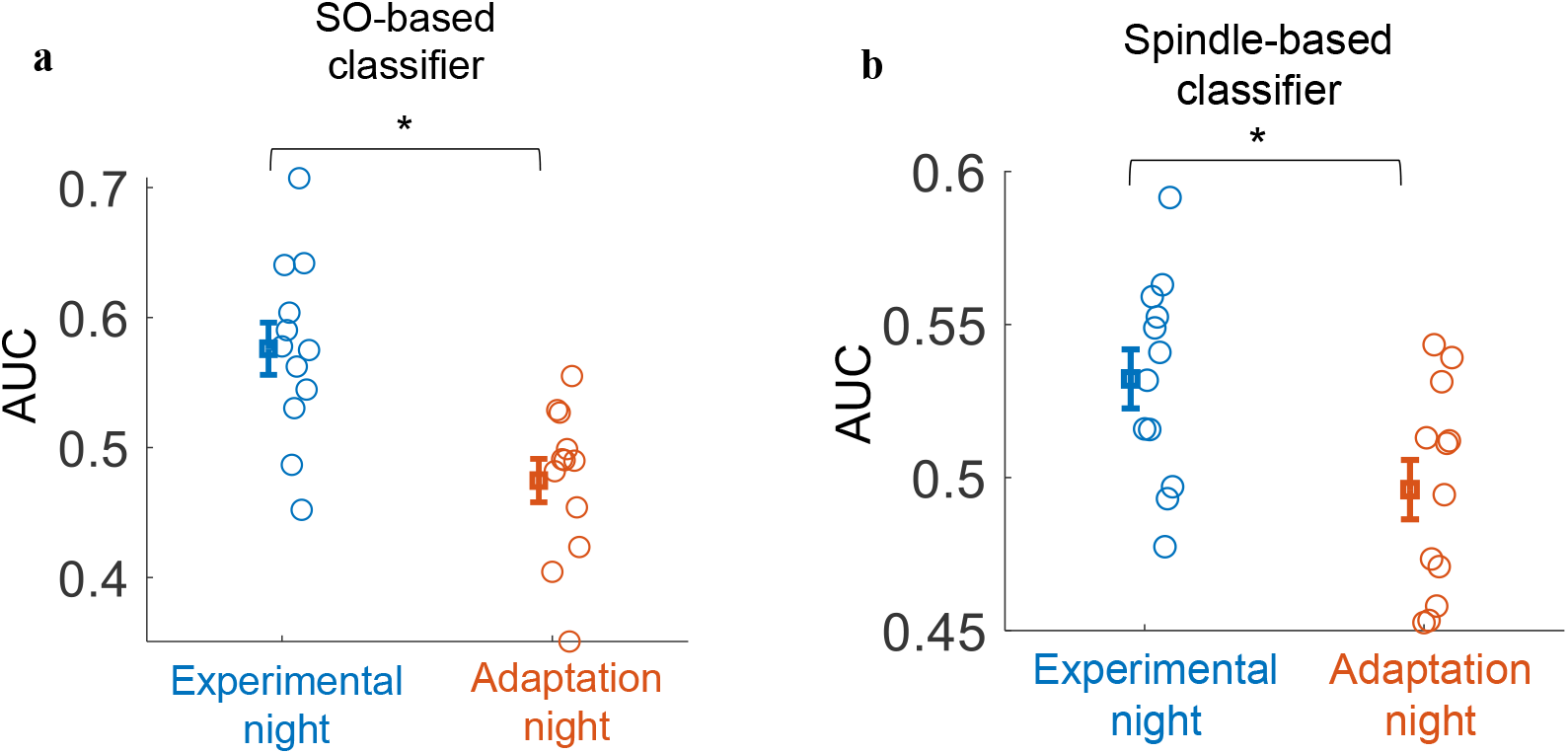
Predicting reactivation using pre-cue features. **a** Classification results of the SO based classifier for the experimental vs the adaptation night for early reactivation (Wilcoxon signed rank test, n=12, P = 0.015, Z = 2.43). b Classification results of the spindle-based classifier for the experimental vs the adaptation night for late reactivation (Wilcoxon signed rank test, n=12, P = 0.028, Z = 2.2).

In addition to the ongoing pattern of SO oscillations, we were interested in how spindles might impact upon the ability of TMR to elicit classifiable reactivations. We therefore repeated the above analysis, but now using spindle features not SOs. We thus trained a spindle-based classifier to predict whether we could use these higher frequency oscillations (11 to 16 Hz), to determine whether TMR would produce a correct classification, using features from channels around the motor area (C5, CP3, C6, and CP4). We thus extracted a binary value representing whether there was a spindle in the 1.5 seconds duration pre-cue (0: no spindle, 1: has spindle), and used this in a decision trees classifier, see methods for details. This showed that we can discriminate between correctly classified and incorrectly classified trials, only in the experimental night and not the adaptation night (Wilcoxon signed rank test for experimental vs. adaptation, n = 12, p = 0.028, z = 2.2), Fig. 3b. Subsequently, we analysed the trials of each participant to determine whether it was the presence or absence of spindles that predicted the correct classification by the reactivation classifier. This showed that trials with fewer pre-cue spindles are more likely to have late reactivation (Supplementary Fig. 3). This is in keeping with a study which showed that significant post-cue reactivation was observed in trials with low pre-cue sigma power^11^. Spindles have been shown to have a periodicity of about 4 seconds^40^, thus, it is possible that the occurrence of pre-cue spindles which prevented post-cue spindles and reactivation in that study^40^ also prevented late reactivation in our study. However, there was no such relationship with early reactivation. Overall, these results suggest that we can use spindle features to predict when to deliver TMR in order to trigger a classifiable reactivation.

### Characteristics of detected reactivations

Because this is a motor task, we wanted to know whether classification of reactivation was derived from the channels over the motor area. We therefore applied the reactivation classification again using only motor channels (C5, CP3, P7, C6, CP4, and P8), both for training in wake and testing in sleep. This motor channel only analysis showed a very similar classification pattern to the results from our all-electrode analysis (Spearman r = 0.99, p < 0.0001, for both peaks when we correlate the classification performance across participants), suggesting that the all-electrode analysis effect shown in Fig. 2a is actually driven by motor electrodes.

Because sleep is characterised by relatively low frequencies such as SOs (0.5 – 1.5 Hz), delta waves (1.5 – 4 Hz), and theta (4 – 8 Hz), we hypothesised that these would be the most important for our classification. To investigate this, we applied a low pass filter with cut-off frequency of 10 Hz. The resulting classification pattern was similar to the result without this filter in Fig. 2a (Between-analyses Spearman on classification: r = 0.93, p < 0.0001 for early peak and r = 0.96, p < 0.0001 for late peak), suggesting that the low frequency pattern is important for classification.

As shown in Fig. 2a, we found that reactivation could occur at either of the two different timepoints - either early after the onset of the cue, or approximately one second later. This demonstrates the temporal characteristics of reactivations within trial duration. We also wanted to examine the characteristics of reactivations occurring at these two different times across the time course of stimulation. Our prior work on this task suggested that classification performance decreases as the number of stimulations in a night increases^41^. We tested whether more correct classifications occur before or after the middle of the stimulation time. Thus, we indexed the correctly classified trials for early/late reactivation to range from 0 (first trial in stimulation) to 1 (last trial) for every participant. We then compared the indices to 0.5 (middle of stimulation) across participants. This revealed that reactivations could be detected to a similar extent at any time during stimulation and was not more prevalent at the beginning or end of stimulation time. Neither reactivations which occurred right after the TMR tone, nor reactivations which occurred ~1 second after the TMR tone, differed significantly from the middle of the stimulation time (Wilcoxon signed rank test, n = 12, p = 0.39, z = 0.86, and p = 0.58, z = 0.55 for early and late reactivations, respectively).

Finally, we wanted to examine how the performance of early and late reactivations varied across the night of stimulation. Thus, we obtained a performance curve across stimulation time for both early and late peaks by observing the changes of classification performance during the time of that peak throughout trials of stimulation (Supplementary Fig. 1). We used a 50-trial block to calculate classification performance, and slid this forward by one trial, to progress along the stimulation time. We then normalised the stimulation time to have the range (0 to 1), with 0 being the first stimulation in the night and 1 the last stimulation. Interestingly, classification performance between the two peaks differed around approximately 0.6, that is, at 60% of the way through stimulation time, with early reactivation more likely to occur at this time (Supplementary Fig. 1).

### The relationship between behaviour and classification performance

Some prior reports have shown a positive relationship between detectable reactivation after TMR tones and the extent of TMR related behavioural benefit^9,10,42^. We searched for this relationship in our data, by testing for correlations between classification and behavioural performance. Because different trials classified correctly at early and late timepoints after the cue, and because such temporally distinct reactivation may potentially also have distinct functional characteristics, we performed correlations twice, using the classification rate at the early peak and then at the late peak. This revealed a negative correlation between pre-sleep reaction time for the reactivated sequence, and classification AUC in the early peak (Spearman r = −0.60, uncorrected p = 0.04), Fig. 4a. In other words, faster pre-sleep performance was associated with a higher classification rate immediately after the TMR cue. This could index a stronger representation that could reactivate more easily, or in a more classifiable form.

**Fig. 4.**
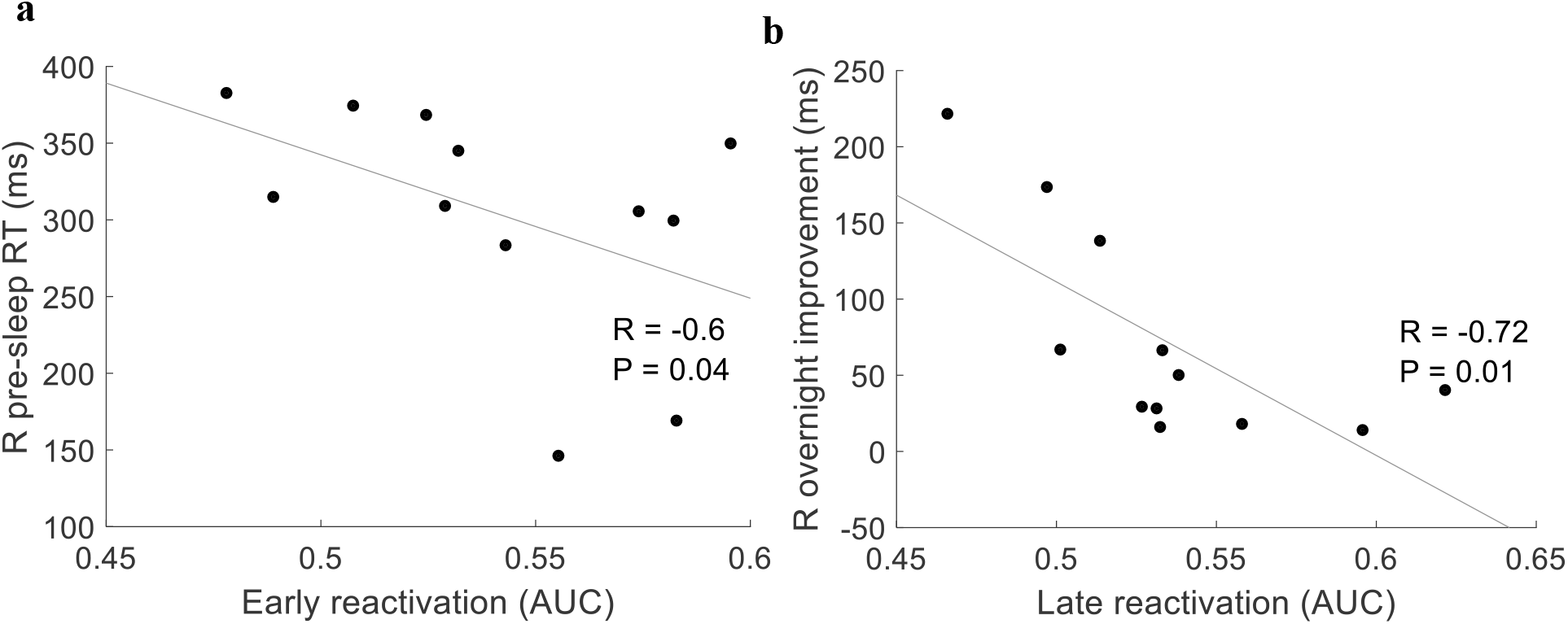
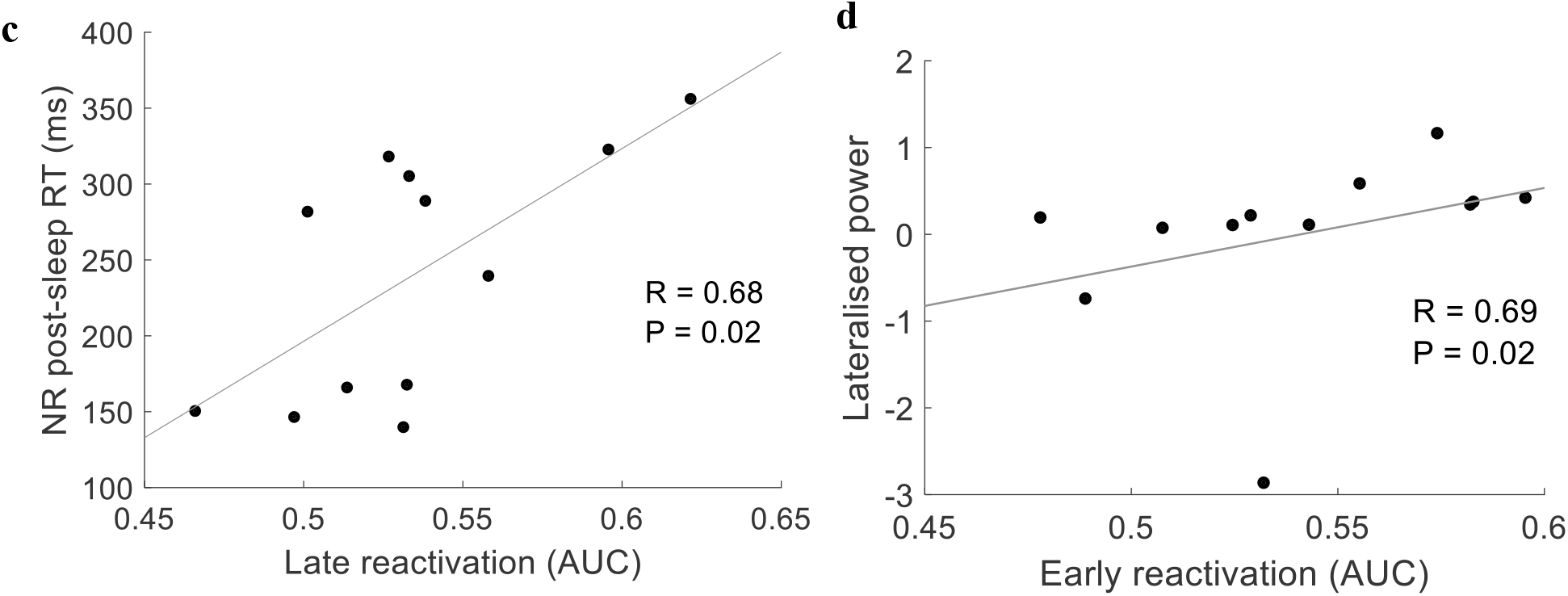
Correlation of classification performance with behavioural results and sigma power. **a** Negative correlation between the classification performance (AUC) of the first peak and the average reaction time (RT) of the last four blocks before sleep for the reactivated (R) sequence (spearman correlation = −0.60, uncorrected p=0.04). **b** Late peak correlated negatively with the overnight improvement of the reactivated sequence (spearman correlation = −0.72, uncorrected p=0.01). c Late peak predicted slower reaction times after sleep for the non-reactivated (NR) sequence (spearman correlation = 0.68, uncorrected p=0.019). **d** Correlation of lateralized sigma power (11 to 16 Hz) (z-transformed) with classification performance for the early peak (Spearman correlation = 0.69, p = 0.016).

Interestingly, the late peak showed quite different associations from the early peak. Here, classification AUC negatively predicted the extent to which responses on the cued sequence sped up across the night of sleep (performance just before sleep – performance early post-sleep), (Spearman r = −0.72, uncorrected p = 0.01), Fig. 4b. We refer to this as overnight improvement, however note that this change is related to reaction time and not improvement in sequence learning, since our measure of sequence learning involves comparison to the random sequence and this measure does not. The AUC of the late peak also predicted slower reaction times for the non-cued sequence after sleep (Spearman r = 0.68, uncorrected p = 0.02), Fig. 4c. Thus, the stronger the late peak, the more slowly the non-reactivated sequence was performed immediately after sleep. These results could suggest that when reactivation occurs late (~1 second after the TMR cue), it somehow disrupts both spontaneous and cued consolidation of the task for both the non-reactivated and reactivated sequence. The idea that late reactivation could have this disruptive property is in-line with a study showing a negative correlation between reactivation and improvement^12^.

### The relationship between sleep spindles and classification performance

Sleep spindles have been strongly linked with memory reactivation^1,43,44^. Work in rodents shows that replays correlate with spindles^45^. Lateralised spindle density during cue periods has been shown to predict TMR-related benefit^23^. We tested for a relationship between sigma power at (11 to 16 Hz) and classification performance. This showed that, even though participants used both hands in this task, right lateralized sigma power, calculated as [Lateralized power = power right area – power left area], was positively associated with the early classification peak, (Spearman r = 0.69, p = 0.016) as shown in Fig. 4d. Thus, the stronger the right lateralised spindle response, the more likely we were to classify reactivation immediately after the TMR cue. This is particularly interesting in light of a prior analysis of our behavioural data showing TMR-related improvement in the weaker left, but not the stronger right hand over sleep^26^. This correlation suggests that a lateralised spindle response may provide a marker for more classifiable early reactivation.

## Discussion

This study shows that TMR cues are more likely to result in classifiable reactivation when applied during the up-going phase of the SOs. We also show that the pattern of ongoing SOs and spindles before a TMR cue can be used to predict whether each cue will produce a classifiable reactivation. Importantly, the resultant reactivations did not reoccur after the TMR cue, instead occurring either early or late. These findings markedly deepen our understanding of neural reactivations after TMR cues in sleep and may lead to improved methods for efficient boosting of memory via the TMR manipulation.

### Timing of reactivation after the cue

The delay between TMR onset and triggered reactivation, is a matter of current interest. Rodent work showing a rapid reverberation of reactivation between cortex and hippocampus at the millisecond scale has led to the idea that replays may ‘echo back’ again and again after TMR^46^. Other work in rodents^42^ suggests that TMR cued replay can continue to repeat for up to 10 seconds after the offset of the auditory cue and a second cue can interrupt this replay. In humans, a study showed evidence of reactivation about two seconds after TMR, with a suggestion of earlier reactivation immediately after the cue^9^. Another study showed recurrent reactivation after TMR, with the response occurring both immediately after the cue and about two seconds later^10^. Our findings are in keeping with this work, since they suggest that reactivation can occur either immediately after the cue or around one second later. Because our inter-trial interval was only 1,500 ms, it is possible that reactivations could occur even later, but the next TMR cue would likely have prevented this in our current design. Notably, although we saw no evidence of recurrent reactivation in the timescale of seconds, this does not rule out the idea of extremely rapid recurrence of reactivation at the millisecond timescale, as suggested in^46^.

Since our data show that reactivations can be identified either early or late after the cue, we must ask whether such differences in timing are functionally important. Early reactivation was positively related to pre-sleep behavioural performance, while late reactivation was instead negatively related to the extent of overnight improvement. It is difficult to interpret these findings, but one possible explanation could be that a strongly encoded memory of the task leads to more immediate reactivation after a TMR cue while a weaker memory trace leads to late reactivation which might actually disrupt consolidation of the task if memories become distorted during the delay.

### Optimal timing of TMR cues

The exact mechanisms by which TMR triggers reactivation are unknown, but the up-going phase of the SO is more reactive to stimulation than the down-going phase, after all, neurones are preparing to fire as the oscillation approaches its peak and beginning a silent period as it enters the trough. Auditory stimulation after the negative peak of the SO, during the up-going phase, has been shown to produce a higher amplitude ERP than stimulating during the down-going phase^16^. TMR can therefore also be expected to have different impact in the up-vs. down-going phase of the SO.

Cortical SOs, thalamo-cortical spindles and sharp wave ripples are thought to be key for memory consolidation, with the up-going SO driving spindle-ripple events with reactivation^13,47,48^. TMR to the up-going phase of the SO has been shown to improve memory^14,15^. This relationship between SO phase and reactivation is clear in the current work, as we show that stimulating the SO up-going phase maximises detectability of reactivation.

SOs are highly heterogeneous, differing both in locus of generation and shape. For instance, SOs differ in period, trough depth, and peak to trough slope^19,22^. Importantly, the SO down-state is thought to be required for the generation of a thalamic down-state which triggers a spindle^49^. Given the association between memory reactivation and spindles, along with the fact that spindle initiation requires a sharp SO trough, it is reasonable to suppose that TMR stimulation of some SOs may be more likely to trigger reactivation than TMR stimulation of others. For instance, SOs with a deeper trough or steeper slope, or some combination of these might be more likely to carry reactivation-bearing spindles. Such differences could explain why we were able to predict which stimulations would be successful based on the features of the ongoing SO before the TMR cue, although, notably, the combination of features was necessary and no single SO feature was sufficient for this prediction.

We also found that trials with fewer pre-cue spindles are more likely to have late reactivation (Supplementary Fig. 3). This is in good keeping with work from Wang and colleagues showing that lower pre-cue sigma power predicted more post-cue reactivation, and that such reactivation begins around one second after the onset of the cue^11^. Such predictive analysis could potentially be used to boost the efficacy of TMR by ensuring that stimulation occurs only at the times when it is most likely to be effective. This could minimise any potential disturbance from TMR, which does often lead to arousals when delivered indiscriminately. Such increased precision of cue delivery could be important for translation of the TMR technique from lab to the home environment.

### Control analyses

Our control for detecting memory reactivation was two-fold. Firstly, we encapsulate the identity of a memory using its EEG pattern during wake and use this as a guidance for detecting the reactivation of this memory in sleep after TMR cues. We achieve this by training our classification models on EEG from wake during wakeful encoding and testing them in sleep. This procedure ensures that classification strength is caused by the reinstatement of the same encoded memories and related to genuine re-processing of memories during sleep. Our work builds on paradigms showing the discriminability of cued categories in sleep data without the inclusion of wake^9,50^, as well as an approach that includes only the features that caused category discrimination in wake^11^. We also use a control night to ensure that our classification results of the experimental night are not caused by sound induced noise in EEG.

### Summary

In this study, we show that reactivations can occur either early or late after a sound cue, and appear to have different functional significance depending on this timing. We also show that TMR delivered during the SO up-going transition is more likely to be associated with classifiable reactivation, probably because it heralds the spindle-bearing upstate. Finally, we show that both pre-cue SO morphology and spindle incidence can be used to predict TMR cued reactivation. This method will allow more efficient TMR stimulation, paving the way for the development of wearable devices to effectively stimulate memory consolidation in the home environment.

## Methods

### Participants

The current study uses EEG from human participants (n=15), mean age: 23.4 years, 7 females. Participants spent an adaptation night in the lab, then in the experimental night, they completed a SRTT before and after sleep. All participants were right-handed and none of them reported familiarity with SRTT. All participants had normal or corrected-to-normal vision, normal hearing, and no history of physical, psychological, neurological, or sleep disorders. Responses in a pre-screening questionnaire reported no stressful events and no travel before commencing the study. Participants did not consume alcohol in the 12 hours before the study and caffeine in the 24 hours prior to the study or perform any extreme physical exercise or nap. This study was approved by the School of Psychology, Cardiff University Research Ethics Committee, and all participants gave written informed consents^26^.

### Experimental design

Participants were asked to sleep in the lab before doing the SRTT training. During this control night, sounds were played with the same criteria as the actual experiment, but importantly had not yet been associated with any task. During SWS, TMR cues of the cued sequence (12-items) were presented to participants with a 20-second pause between the presentation of each sequence.

Participants performed a SRTT (SRTT; adapted from^23^. Sounds cued four different finger presses. We delivered the sound cues during SWS. Participants learned two 12-item sequences, A and B, A: 1 2 1 4 2 3 4 1 3 2 4 3 and B: 2 4 3 2 3 1 4 2 3 1 4 1. The location indicated which key on the keyboard needed to be pressed as quickly and accurately as possible: 1 – top left corner = left shift key; 2 – bottom left corner = left Ctrl; 3 – top right corner = up arrow; 4 – bottom right corner = down arrow. Sequences had been matched for learning difficulty; both contained each item three times. The blocks were interleaved so that a block of the same sequence was presented no more than twice in a row, and each block contained three repetitions of a sequence. There were 24 blocks of each sequence (48 blocks in total), and each block was followed by a pause of 15 seconds wherein feedback on reaction time (RT) and error-rate were presented. The pause could be extended by the participants if they wanted. After the 48 blocks of sequences A and B, participants performed four more blocks that contained random sequences. They contained the same visual stimuli and an ‘R’ displayed on the centre of the screen. Two of these blocks were paired with the tone group of one sequence (reactivated in sleep), and the other two were paired with the tone group of the other sequence (non-reactivated). Participants were aware that there were two twelve-item sequences and each sequence was indicated with ‘A’ or ‘B’ appearing on the centre of the screen, but they were not asked to learn the sequences explicitly. Counterbalancing across participants determined whether sequence A or B was the first block, and which of the sequences was reactivated during sleep.

Each sequence was paired with a group of pure musical tones, either low tones within the 4th octave (C/D/E/F) or high tones within the 5th octave (A/B/C#/D). These tone groups were counterbalanced across sequences. For each trial, a 200ms tone was played, and at the same time a visual cue appeared in one of the corners of the screen. Participants were instructed to keep individual fingers of their left and right hand on the left and right response keys, respectively. Visual cues were neutral objects or faces, used in previous study^23^, which appeared in the same position for each sequence (1 = male face, 2 = lamp, 3 = female face, 4 = water tap). The nature of the cues (objects/faces), participants were told, was irrelevant. Visual cues stayed on the screen until the correct key was pressed, after which an 880 ms inter-trial interval followed.

After completion of the SRTT, participants were asked to do the same task again, but were instructed to only imagine pressing the buttons. This imagery task consisted of 30 interleaved blocks (15 of each sequence), presented in the same order as during the SRTT. Again, each trial consisted of a 200 ms tone and a visual stimulus, the latter being shown for 270 ms and followed by an 880 ms inter-trial interval. There were no random blocks during the imagery task and no performance feedback was presented during the pause between blocks.

In the morning following the experimental night, participants were asked to perform the task again, motor imagery first, then SRTT. Eventually, they were asked if they remember the images locations of the two sequences, to see if one sequence is recalled better than the other one. Motor imagery data set of each participant was used for classification. The adaptation/control night was useful for eliminating the possibility that a classifier could merely classify sound induced effects on the EEG. Thus, if the classifier can classify the experimental night but not the adaptation night this suggests that it is classifying memory reactivations.

### Recording and pre-processing

Data were extracted using 21 electrodes, following the 10-20 EEG system. 13 of the electrodes were placed on standard locations, namely: FZ, CZ, PZ, F3, F4, C5, CP3, C6, CP4, P7, P8, O1, and O2, and they were referenced to the mean of the left and right mastoid electrodes. Other electrodes were the left and right EOG, three EMG electrodes, which were placed on the chin, and the ground electrode placed on the forehead. The impedance was <5 kΩ for each scalp electrode, and <10 kΩ for each face electrode. Recordings were made with an Embla N7000 amplifier and RemLogic 1.1 PSG Software (Natus Medical Incorporated). PSG recordings were scored by two trained sleep scorers and only the parts scored as SWS were kept for further analyses. Data were collected at 200 Hz sampling rate.

EEG signals were band pass filtered in the frequency range from (0.1 to 30 Hz). Subsequently, trials were cleaned based on statistical measures consisting of variance and mean. Trials were segmented −0.5 to 3 seconds relative to the onset of the cue. Trials falling two standard deviations higher than the mean were considered outliers and rejected if they show to be outliers for more than 25% of the channels. If trials were bad in less than 25% of the channels, they were interpolated using triangulation of neighbouring channels. Thus, 11.7% and 11.9% of trials were considered outliers and removed from the experimental night data and the adaptation night, respectively. Analyses were done using FieldTrip^51^ and Matlab 2018a.

### Wake-to-wake classification and determining a time of high discrimination

We started the analysis by performing a wake-to-wake motor imagery classification. This was performed for each participant separately, with trials serving as observations, labelled according to the hand they belong to. EEG signals were pre-processed and features were extracted by calculating time-domain amplitude averages of 80 ms (40 ms before and 40 ms after every time point). Subsequently, features were fed to a linear discriminant analysis (LDA) classifier^52^. Training and testing was done on wake data in a time x time fashion^53^. The classifier was trained on a specific time point and tested with all time points using a 5-fold cross-validation to build one row in the time x time classification, illustrated in Fig. 5a. We assumed that if classification performance is not at a considerably high rate during wake then this would decrease the possibility of classifying sleep reactivation where noise is higher. Consequently, we chose the participants that had wake-to-wake classification performance with area under the ROC curve (AUC) >= 0.7, (n = 13). One participant was neglected because of a technical problem during the collection of sleep data. The rest of data was used for classification (n = 12). We also do realise the rich literature of motor imagery classification with common spatial patterns (CSP) and other methods^54–58^. However, given the differences between wake and sleep data sets and their different nature of noise and oscillations we decided to directly use time domain features with our classifiers.

**Fig. 5.**
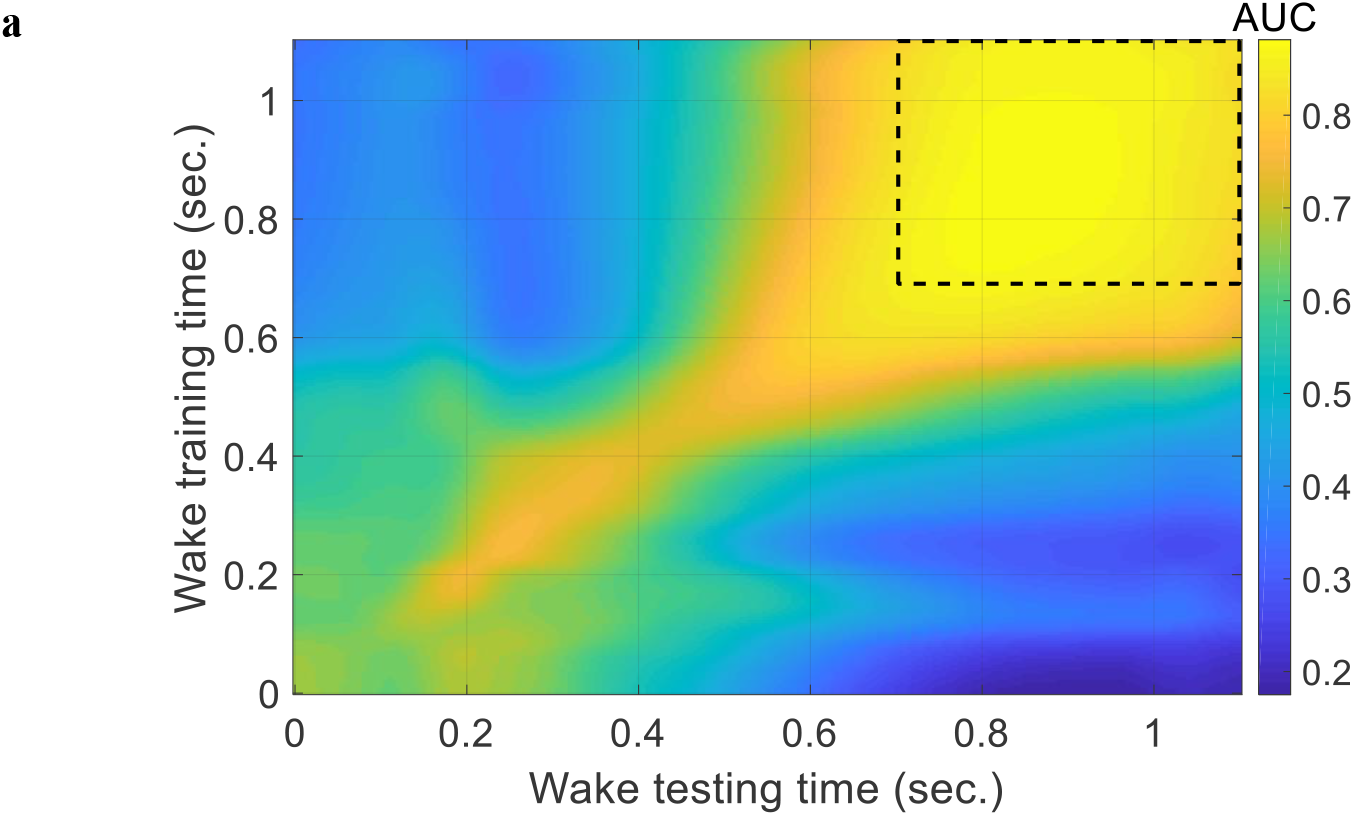
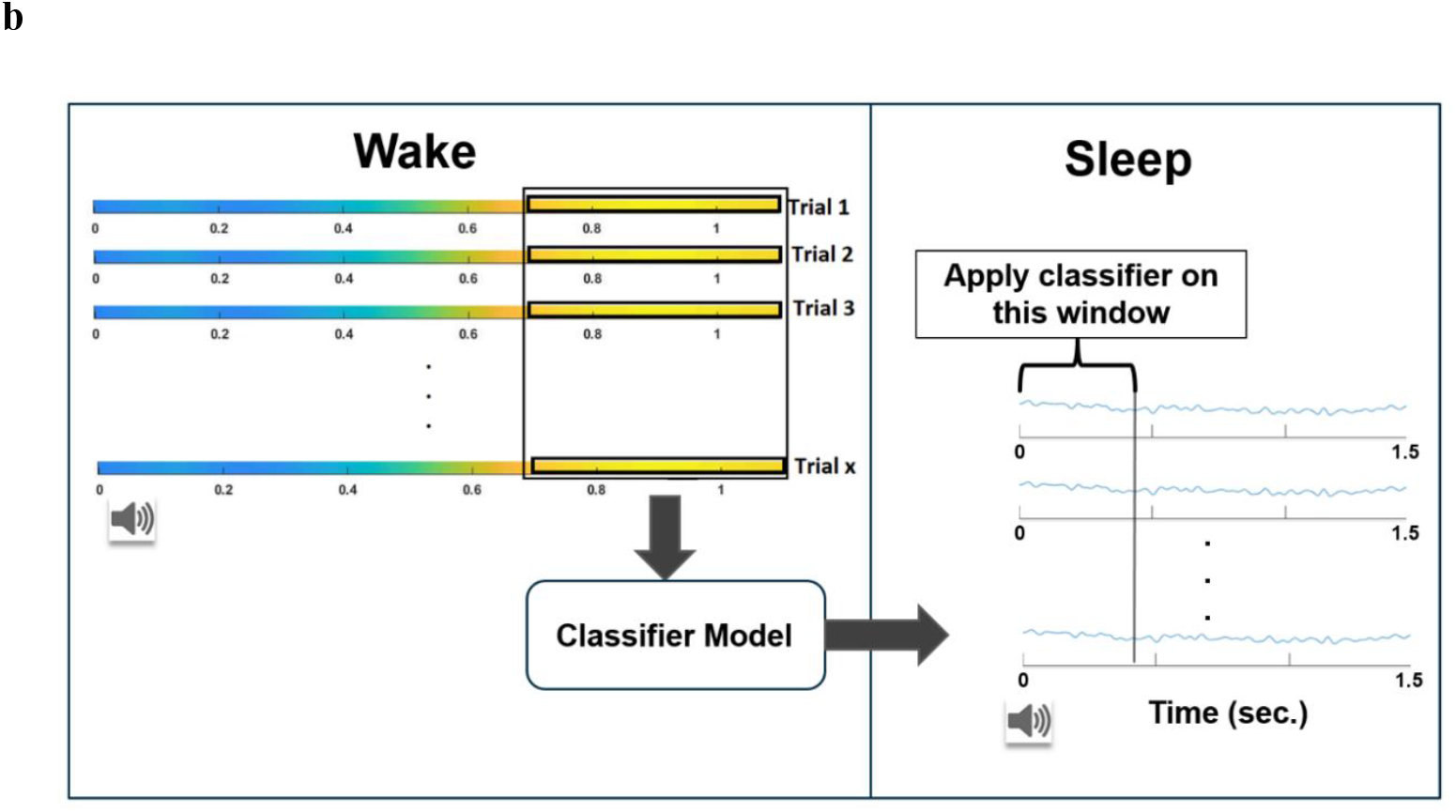
Classification of left-vs. right-hand. **a** Grand average classification AUC for left-vs. right-hand motor imagery using a sliding 80 ms smoothing window and LDA classifiers, dashed box represents the time of interest (TOI). b Illustration of classification procedure (training with wake and testing with sleep) which is applied for both the experimental night and adaptation (control) night. A sliding window approach is used, wherein a classifier is tested on a window from sleep and the classification result replaces the centre of that window then the window is slid by one time point to construct a performance curve across trial time in sleep.

Initial investigations revealed a higher classification performance for left-vs right-hand (where both fingers were aggregated into one class) than for faces vs objects. Therefore, we conducted the analysis on right-vs left-hand imagery. The trial length was defined as the duration between cue onsets (1.1 seconds in wake). Sound cues had a duration of 200 ms and were played from time 0 of trials. During sleep, trial length was 1.5 seconds.

Classification of motor imagery during wake shows a time period with maximum classification performance (marked with dashed box in Fig. 5a). This time region should be useful for discriminating left hand and right hand. We defined this time period as the time of interest (TOI). A TOI is a time window that has high classification rate, indicating its ability to discriminate the classes. It acts as a temporal marker of expected discrimination. To locate this window, we used a threshold of 0.85 on the average classification AUC from all participants.

### Wake-sleep classification

Once we had built a classifier on wake data, we tested it on sleep data. We applied it to sleep using a sliding window approach, as shown in Fig. 5b. Using the sliding window approach the classification was applied on the first testing window in sleep, for example: [0 to 0.38] second, which matches the length of the TOI. Then, the classification performance is placed at the centre time of this window, i.e., at 0.190 second. Subsequently, the sliding sleep window is shifted by one time point, and the process is repeated. Thus, the results of classification are AUC values across time.

The wake-to-sleep classifiers used the concatenated averages inside the TOI as features. These concatenated time points were reduced to the most informative contiguous time points using mutual information on wake data for each participant. We reduce the features to the most informative time points since the reactivation might be temporally short compared to wake activation. Consequently, we slide a shorter window that contains the most informative features which enables the classifier to detect the reactivation if it was temporally short or long. The most informative time points were chosen such that the time points are contiguous and contain the highest 10% of the mutual information values.

We devised a method for removing noisy trials with no TMR effect. We assume noisy trials belong to a new ‘no effect’ class which does not contain discriminative features for right-vs left-hand. The feature values of those trials should fall near the decision boundary in the feature space, in a region where the classifier is uncertain (corresponding to a maximum posterior probability close to 0.5); see Supplementary Fig. 2a, for a representation using two features. Thus, we define trials as ‘no effect’ if they fall in that area. We rejected noisy trials falling in the area of uncertainty and used three hundred clean trials from every participant, those clean trials correspond to a certainty average of 0.86, with 0.1 standard deviation. Importantly, to avoid any bias, this cleaning process was unsupervised, meaning that the information of the ground truth class labels of sleep data was not used. Moreover, we verified that this cleaning process would not be improve classification performance if the data we were trying to clean was random and contained no discriminative information, as illustrated in Supplementary Fig. 2b, which was the case with the control night. It would also not improve classification performance if sleep data was not scattered in the feature space in a similar way to wake samples because the decision boundary position and orientation which are determined using wake will then be meaningless for sleep samples. Thus, this cleaning process only works if sleep data contains discriminative information. Importantly, the exact same cleaning procedure was performed for both the experimental and adaptation night for completeness.

### Preferred TMR phase analysis

Phase information was extracted using Hilbert transform on the band pass filtered signal (0.5 to 2 Hz) using FZ electrode. We divided phase values into two ranges: [0 to π] and (π to 2π], indicating the two transitions: down-going and up-going, respectively. For each participant, we determined the number of correctly classified trials in which TMR fell on either phase range in each night, then normalised by the total number of correct trials. Trials were deemed ‘correct’ when the prediction of the classifier was the same as the respective trial category. We compared the proportion of correct trials where TMR occurred in the down-going and up-going transitions of the SO. The same process was repeated for the incorrect trials of the experimental night also, the correct and incorrect trials of the adaptation night.

### Lateralised sleep sigma power analysis

The lateralized sigma power was calculated using Hilbert transform on the band pass filtered signals (11 to 16 Hz), during the duration: (0 to 0.5) second relative to cue onset which is around the early reactivation. Lateralized spindle power was calculated as: Lateralized power = power right area (C6+CP4) – power left area (C5+CP3)

We tested the correlation between the classification performance of the low-pass filtered signals with cut-off frequency of 10 Hz to be sure that the sigma band is not included. We did this to ensure that the two conditions: lateralised sigma power and classification performance, that we test for correlation are not the same. Classification performance from the low-pass filtered signals still correlated with lateralised sigma power (spearman correlation = 0.61, p = 0.040).

### Reoccurrence of reactivation

We statistically tested if one reactivation (early or late) is more likely to happen or whether reactivation is reoccurring after a sound cue. Thus, we took the accuracy for recurring reactivation (i.e., the ratio of correct trials during the time of both early and late reactivation simultaneously) and compared it to the probability of both reactivations happening simultaneously after a sound cue (the accuracy for early reactivation multiplied by the accuracy for late reactivation) as a chance level. We performed this analysis for every participant and compared the accuracy of reoccurring reactivation to chance level.

### SO based classification

The SO based classification consisted of 200 decision trees ensemble. Leave one out classification was used, wherein the data of all participants except one is used to train the classifier and the left-out participant is used for testing the classifier. This gives a classification result for the left-out participant and the process is then repeated until the classification performance is calculated for all participants. Every decision tree is trained on a random subset of trials from the training set and tested on the testing set and the final result is the aggregated votes from all decision trees. The extraction of SO and spindle events followed the implementation in^59^. Briefly, for spindles, signals were filtered using two-pass bandpass FIR filter between 11 and 17 Hz, subsequently, the root mean square (RMS) power using a window of 200ms. A threshold on power of 86.639 (equivalent to 1.5 standard deviation from the mean of a normal distribution) was applied and spindles were defined as the parts that exceeded that threshold and had duration >300 ms and < 3 seconds. For SO, two-pass FIR bandpass filtering was applied to the signals between 0.3 and 3 Hz. SOs were then extracted as parts with negative deflections that had consecutive zero crossings between 0.25 and 1 second.

### Statistical testing

To assess the statistical significance of the classification results, we compared the classification performance of the experimental night against the adaptation/control night. Sounds played during the adaptation night were the same sounds used in the experimental night but because the adaptation night was before participants had been trained on the experimental task, these sounds were not yet associated with any memories. This control was used to make sure that classification is not derived due to sound induced features/noise in EEG.

Statistical analysis was performed using the classification results of the two nights with cluster based permutation using FieldTrip^51^. Monte Carlo was used with a sample-specific test statistic threshold of 0.05, permutation test threshold for clusters of 0.05, and 10,000 permutations. The correction window used in the test was the whole length of sleep trial, i.e., [0 to 1.5] seconds.

## Data availability

All relevant data generated or analysed are available from the authors upon reasonable request, including the EEG data, behavioural files, all the analyses with MATLAB scripts. Participants private identifications are all anonymized.

## Acknowledgements

This work was supported by the ERC grant SolutionSleep to P.A.L and ERC funded the Ph.D. of M.E.A.A. We would like to thank Miguel Navarrete for his help in the extraction of slow oscillation features for the slow oscillation-based classification and also Lorena Santamaria, Martyna Rakowska and members of our group NaPs for the useful advice.

## Author contributions

A.C.M.K. and P.A.L. designed the experiment. A.C.M.K collected the data. M.E.A.A analysed the data. M.E.A.A., P.A.L., A.C.M.K and M.S.T wrote the manuscript.

## Additional information

## Supplementary Information

is available online.

