## Supplementary figures for "Targeting targeted memory reactivation: characteristics of cued reactivation in sleep"

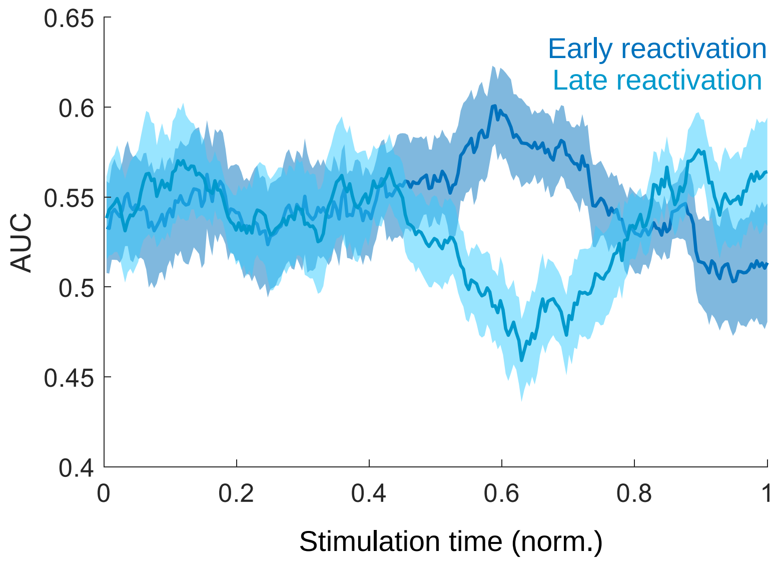


**Supplementary Fig. 1** Performance of classification peaks throughout stimulation time (stimulation time is normalised to have the range (0 to 1)) the performance was calculated for each 50-trial block, the shaded area represents the standard error (SE) and the solid line represents the mean, after the middle of stimulation time, early and late reactivations show different behaviour.

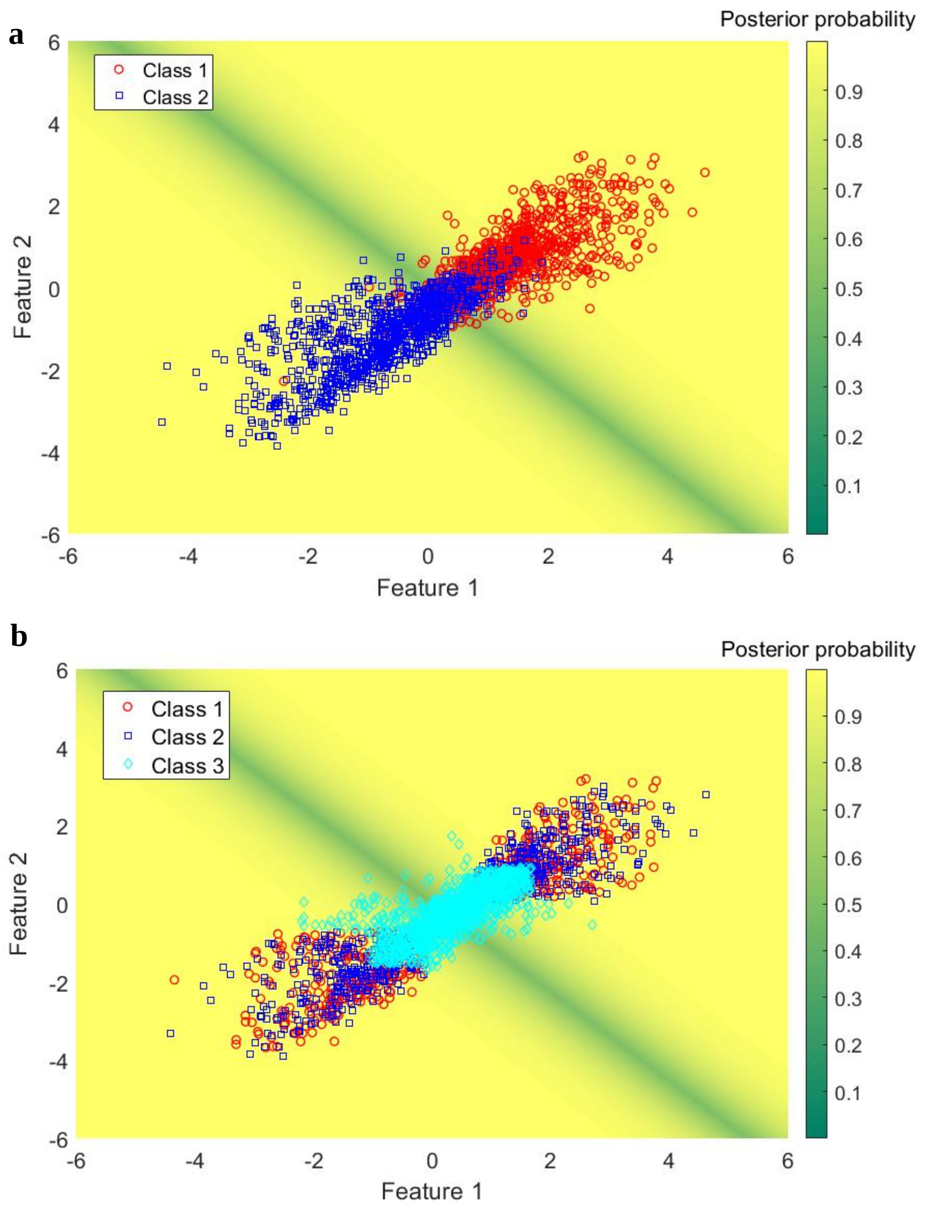


**Supplementary Fig. 2** Illustrating classification feature space using two features from one participant. **a** Classification of left hand (class 1) vs. right hand (class 2), maximum posterior probability among the two classes for every sample is illustrated. Wake samples are shown and the area of 'no effect' is located near the decision boundary and corresponds to low posterior probability (green). **b** If trials were random and do not contain discriminative information then rejecting some trials that fell near the decision boundary (cyan) will not lead to improved classification performance. Thus, if data is random then this cleaning will not improve classification performance.

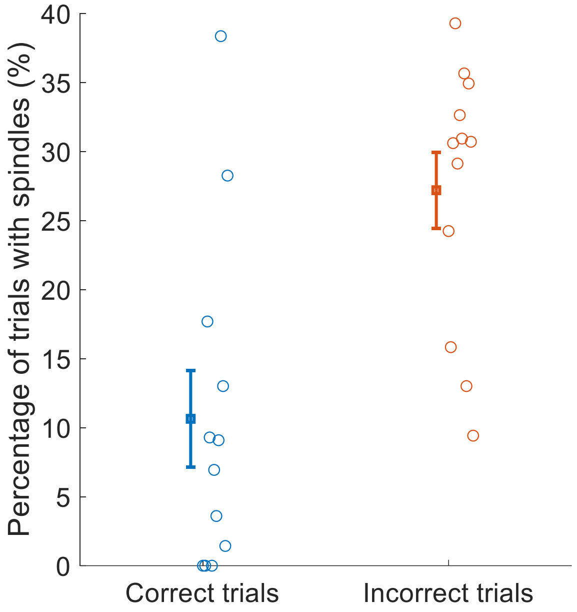


Supplementary Fig. 3 Likelihood of pre-cue spindles (-1.5 to 0) second for correct and incorrect trials. percentages of trials with spindles are shown for correct and incorrect trials of reactivation classifier which shows that the lack of pre-cue spindles is accompanying classifiable post-cue late reactivation. Each circle represents one participant.

| Feature | Description | Variable |
|---|---|---|
| **cosPhase** | Cosine of the phase of auditory stimulation | Continuous |
| **sinPhase** | Sine of the phase of auditory stimulation | Continuous |
| **vSOTrough** | Voltage of SO trough before the click | Continuous |
| **vSOPeak** | Voltage of SO peak in the click wave | Continuous |
| **vSOPeakTrough** | Voltage of SO peak-trough before the click | Continuous |
| **tSONegWave** | Time of duration for the negative wave before click | Continuous |
| **tSOPosWave** | Time of duration for the positive wave before click | Continuous |
| **tSORising** | Time of duration from the trough to zero crossing before click | Continuous |
| **tSOPeakTrough** | Time of duration for the peak to trough before click | Continuous |
| **tStimSOCrossing** | Time between zero-crossing to click time | Continuous |
| **tStimSOTrough** | Time between trough before click to click time | Continuous |

| | | |
|---|---|---|
| **tStimEstimPeak** | Time between click time to wave peak | Continuous |
| **rmsSONegWave** | Area under curve for trough section before click (troughArea) | Continuous |
| **rmsSOPosWave** | Area under curve for peak section before the click | Continuous |
| **rmsSOWave** | Area under curve for all wave previous to stimulation | Continuous |
| **numSOTroughs** | Number of troughs in the negative wave before click | Ordinal |
| **numSOPeaks** | Number of peaks in the positive wave before click | Ordinal |
| **risingSOSlope** | Slope from the trough to zero crossing before click | Continuous |
| **SOwaveRatio** | Duration ratio for the wave before click (Slope) | Continuous |
| **SOhalfWaveRatio** | Duration ratio for the negative wave before click | Continuous |
| **FSonStim** | Presence of fast spindle (11 to 17 Hz) on stimulation | Binary |
| **SSonStim** | Presence of slow spindle (9 to 11 Hz) on stimulation | Binary |
| **existFSonSO** | Presence of fast spindle on the wave before stimulation | Binary |
| **existSSonSO** | Presence of slow spindle on the wave before stimulation | Binary |

Supplementary Table 1 Description of SO features used for predicting reactivation.
